# Can AlphaFold2 predict protein-peptide complex structures accurately?

**DOI:** 10.1101/2021.07.27.453972

**Authors:** Junsu Ko, Juyong Lee

## Abstract

In this preprint, we investigated whether AlphaFold2, AF2, can predict protein-peptide complex structures only with sequence information. We modeled the structures of 203 protein-peptide complexes from the PepBDB DB and 183 from the PepSet. The structures were modeling with concatenated sequences of receptors and peptides via poly-glycine linker. We found that for more than half of the test cases, AF2 predicted the bound structures of peptides with good accuracy, C*α*-RMSD of a peptide < 3.0 Å. For about 40% of cases, the peptide structures were modeled with an accuracy of C*α*-RMSD < 2.0 Å. Our benchmark results clearly show that AF2 has a great potential to be applied to various higher-order structure prediction tasks.

## 1 Introduction

In CASP14, DeepMind introduced the version of their protein structure prediction model, AlphaFold2 (AF2), demonstrating that it can predict the protein structures with remarkable accuracy (Callaway [2020]). The median GDT-HA score of AF2 predictions was 78%, which is comparable to crystal structures (Pereira et al.). This high accuracy indicates that almost all single-domain protein structures can be determined accurately with AF2. In addition, on July-23-2021, DeepMind and EMBL released the model structures of all human proteome and the other 20 organisms. These results will open new possibilities for biology and its related fields.

Based on the achievement of AF2, we tried to investigate whether AF2 can be applied beyond the prediction of single domain structures. As the first attempt, we evaluated AF2’s ability to predict protein-peptide complex structures. Protein-peptide complexes are playing essential roles in biological processes. Around 154∼0% of protein-protein interactions (PPI) are estimated to be involved with protein-peptide interactions (Petsalaki and Russell [2008]). Therefore, the accurate modeling of protein-peptide complex structure is critical in understanding the mechanisms of biological reactions and development of peptide drugs (Ciemny et al. [2018]).

Various algorithms and programs have been developed to predict protein-peptide complex structures (Ciemny et al. [2018]). Protein-peptide modeling approaches can be categorized into three groups: (1) template-based (Lee et al. [2015]), (2) local docking (London et al. [2011]), and (3) global docking (Zhou et al. [2018]). A well-organized summary of protein-peptide modeling methods can be found in the reference (Ciemny et al. [2018]). Template-based methods generate model structures based on similar known complex structures. Thus, the approaches inherently rely on the existence of known structures and cannot be applied to novel targets. Local docking approaches perform peptide docking near a preset binding site provided by a user, which requires prior knowledge on a target protein.

Unlike template-based and local docking approaches, global docking approaches perform complex structure modeling solely based on a receptor structure and a peptide sequence. Thus, global docking approaches are general and can be applied to novel targets. However, due to a large degree of freedom, global docking approaches are computationally challenging and show relatively lower accuracy.

In this preprint, we investigated whether AF2 can predict protein-peptide complex structures only with sequence information. It has been shown that protein-protein complex structures can be modeled using coevolutionary information (Weigt et al. [2009], Schug et al. [2009], Balakrishnan et al. [2011]). Thus, if AF2 has learned how to extract coevolutionary information from multiple sequence alignment and interpret it to structural information succesfully, AF2 would be possible to predict the quaternary structures of proteins. Moriwaki and Ohue showed that AF2 may be able to predict the structures of protein-protein or protein-peptide complexes (Moriwaki, Ohue). In a similar vein, we assess the modeling power of AF2 for protein-peptide complexes by modeling the structures of 386 protein-peptide complexes after concatenating the sequences of receptors and peptides using a poly-glycine linker. Therefore, our benchmark corresponds to the global protein-peptide docking. We found that for more than half of the test cases, AF2 predicted the bound structures of peptides with good accuracy, C*α*-RMSD of a peptide < 3.0 Å. Our test results clearly show that AF2 has a great potential to be applied to various higher-order structure prediction tasks.

## 2 Methods and Materials

### 2.1 Dataset

In this study, we used two sets of protein-peptide complexes. The first set consists of 203 protein-peptide complexes randomly selected from PepBDB (Wen et al. [2019]). We used the PepBDB release-2020-03-18 and applied the following two filtering criteria to select target complexes.

1. The sequence lengths of receptor and peptide should be greater than or equal to 10.
2. Either of the sequence lengths of receptor and peptide should be greater than or equal to 50.
3. There are no non-canonical amino acids.

The second set, PepSet, consists of 183 protein-peptide complexes, which were used for comprehensive benchmarking of protein-peptide docking programs (Weng et al. [2020]). The following filtering criteria were used to compile the set.

1. The peptide length ranges from 5 to 20.
2. The resolution is ≤ 2.0 Å.
3. Peptides do not include non-canonical amino acids.
4. The sequence identity between any two protein monomers interacting directly with peptides is less than 30%.
5. The bound structures have the corresponding unbound structures with a sequence identity > 90%.
6. The RMSD of the backbone atoms of the residues in the bound structures within 10 *AA* from the peptide and the corresponding residues of the unbound structure is less than 2.0 Å.

### 2.2 Modeling protein-peptide complex using AlphaFold2

We performed protein-peptide complex modeling using AF2 (Jumper et al. [2021]) with a sequence that protein and peptide sequences were concatenated with 30 glycine residues. The source code of AF2 was retrieved from the github repository (https://github.com/deepmind/alphafold) on July-16-2021. No template information was used for structure modeling by setting the *max_release_date* parameter as 1979-Jan-01. The *full_db* preset was used for modeling, which is the eight times faster than the *casp14* setting, which was used in CASP14. For all targets, five models were generated.

### 2.3 Evaluation metrics

We evaluated the quality of protein-peptide complex structures using the following three metrics: (1) complex RMSD, RMSD measured by the superposition of the whole protein-peptide complex (All_RMSD), (2) peptide RMSD, RMSD measured by the superposition of the peptide only (Pep_RMSD), and (3) peptide RMSD after the optimal superposition using a receptor structure (SP_RMSD, superposed peptide RMSD). In this study, we considered predictions with SP_RMSD ≤ 2.0 Å are high-quality and with SP_RMSD 4.0 ≤ Å near-native. SP_RMSD is conceptually similar to IL_RMSD used for the comprehensive benchmark (Weng et al. [2020]). All RMSD values were calculated with backbone C*α* atoms using the *SVDSuperimposer* implemented in BioPython (Cock et al. [2009]). No distance cutoff between a receptor and a peptide was considered for RMSD calculations, which may lead to high RMSD values for unsuccessful predictions.

## 3 Results and Discussion

Table 1 shows the statistical summary of modeling quality of model 1 of the PepBDB set. It is noticeable that the median of SP_RMSD is 2.3 Å indicating that more than half of predictions were near-native predictions. The first quartile values of SP_RMSD and All_RMSD are 0.79 and 1.22 Å corresponding to highly accurate structures with atomic details. Among 203 targets, 90 complexes were predicted to be high quality, SP_RMSD < 2.0 Å. Also, in terms of All_RMSD, the median value is 2.61 Å suggesting that the whole complex structure can be predicted with the current protocol accurately. In addition, the high accuracy of the models indicates that AlphaFold can be a tool for predicting peptide-binding regions without any prior information.

**Table 1:** The summary of modeling quality of the PepBDB set

|  | Top |  |  | Best |  |  |
| --- | --- | --- | --- | --- | --- | --- |
|  | SP_RMSD | All_RMSD | Pep_RMSD | SP_RMSD | All_RMSD | Pep_RMSD |
| Q1 | 0.79 | 1.22 | 0.99 | 0.75 | 1.20 | 1.07 |
| Q2 (median) | 2.30 | 2.61 | 2.09 | 1.86 | 2.29 | 1.91 |
| Q3 | 7.56 | 7.05 | 3.74 | 5.44 | 5.83 | 3.48 |
| min | 0.11 | 0.37 | 0.15 | 0.11 | 0.37 | 0.15 |
| max | 50.1 | 24.8 | 12.7 | 30.0 | 22.3 | 12.5 |

The results of the best model among five predictions in terms of All_RMSD are summarized on the right side of Table 1. When the best among five models are considered, the median of SP_RMSD and All_RMSD are 1.86 and 2.29 Å. With a threshold of SP_RMSD < 2.0 Å, 104 targets were predicted accurately among the 203 targets, corresponding to a success rate of 51%.

The modeling results of PepSet are summarized in Table 2. Overall, RMSD values are similar to those of the PepBDB set. The median SP_RMSD of the top models is 2.88 Å, which is slightly worse than that of the PepBDB result. When the best models are considered, the SP_RMSD value decreases to 1.79 Å. In summary, it is clear that AF2 can predict protein-peptide complex structures accurately based on the modeling results of 386 protein-peptide complexes,

**Table 2:** The summary of modeling quality of the best model of PepSet

|  | Top |  |  | Best |  |  |
| --- | --- | --- | --- | --- | --- | --- |
|  | SP_RMSD | All_RMSD | Pep_RMSD | SP_RMSD | All_RMSD | Pep_RMSD |
| Q1 | 0.82 | 1.52 | 0.91 | 0.66 | 1.39 | 0.87 |
| Q2 (median) | 2.88 | 3.63 | 1.92 | 1.79 | 2.45 | 1.94 |
| Q3 | 7.16 | 7.16 | 3.14 | 4.37 | 4.92 | 2.79 |
| min | 0.10 | 0.42 | 0.13 | 0.08 | 0.42 | 0.13 |
| max | 36.0 | 29.6 | 9.1 | 24.4 | 29.6 | 8.60 |

Interestingly, modeling accuracy does not depend on the sequence length of a peptide or a complex. The SP_RMSD values and the peptide and complex sequence lengths of the PepBDB set models are illustrated in Figure 1. It is observed that there is almost no correlation between the model accuracy and sequence lengths. This indicates that AF2 modeling does not rely on the number of contacts between a receptor and a peptide or shape complementarity, which is critical for many physics-based or empirical peptide docking programs. This result shows that the strength of the coevolutionary signal is independent of sequence lengths.

**Figure 1:**
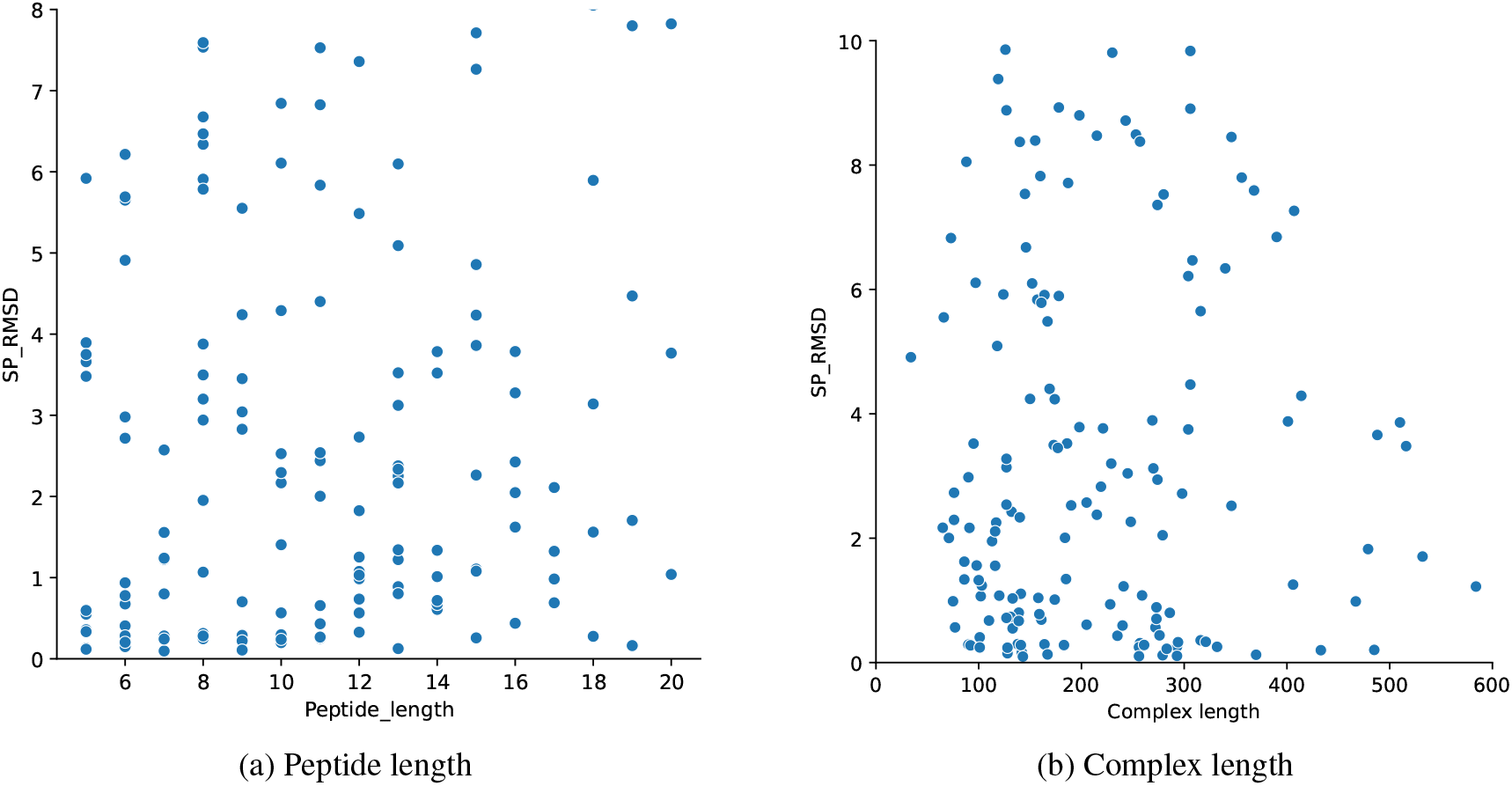
Relationship between model accuracy in SP_RMSD and (a) peptide and (b) complexe sequence lengths

Two successful predictions are illustrated in Figure 2. In the case of 1B2D, the sequence lengths of receptor and peptide are similar. The modeled peptide structure is almost perfectly superposed with the crystal structure. In the case of 6RM8, a long peptide structure is similar to the crystal structure. The helical structure and the following disordered region of the peptide are closely following the crystal. Only the terminal region of the peptide deviates from the crystal.

**Figure 2:**
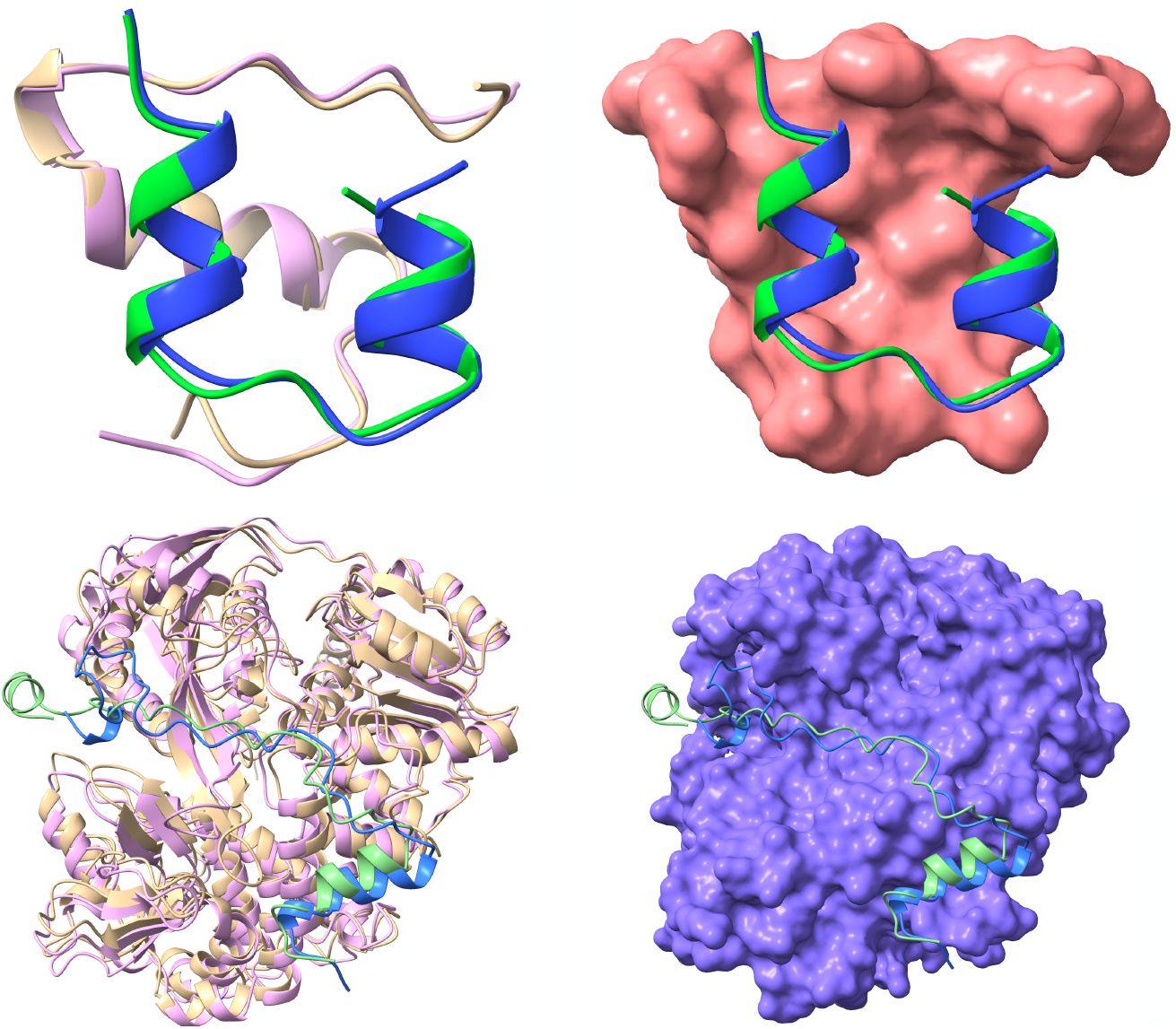
Modeling examples of successful predictions: (top) 1B2D and (bottom) 6RM8. The crystal structures of a receptor and a peptide are colored in gold and green. The model receptor and peptide structures are colored in pink and blue. The glycine linker is not displayed for clarity.

Two unsuccessful prediction results are displayed in Figure 3. In the cases of 4C5A and 5GOW, AF2 could not find proper binding sites. The poly-glycine linker region is fully elongated, and a peptide is predicted to be far away from the receptor. AF2 may have predicted that the given receptor and peptide pairs would not bind.

**Figure 3:**
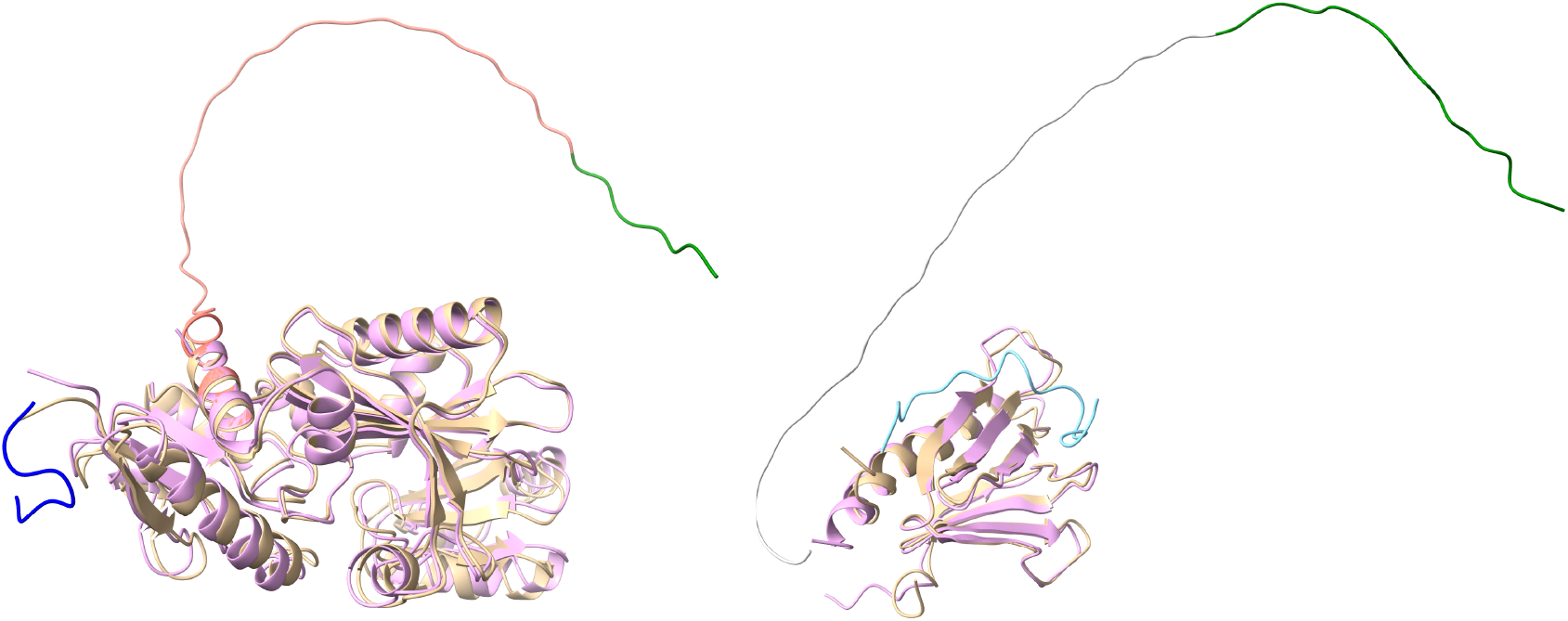
Modeling examples of bad predictions: (left) 4C5A and (right) 5GOW. The crystal structures of a receptor and a peptide are colored in gold and green. The model receptor and peptide structures are colored in pink and blue.

## 4 Conclusion

In this study, we showed that AF2 can predict the protein-peptide complex structures accurately without template information. It is remarkable that AF2 generated accurate model structures using only the concatenated sequence of a receptor and a peptide via a poly-glycine linker. When the top models are considered, 44% and 61% of the PepBDB set targets were predicted with accuracy of SP_RMSD < 2.0 Åand SP_RMSD < 4.0 Å, respectively. This level of accuracy is comparable or better than existing protein-peptide modeling methods without using template information. However, we cannot completely rule out the possibility that AlphaFold may have used protein-peptide complex structures information during training. It is also worth investigating how much model qualities would improve with template information. These results suggest that AF2 learned how to extract interaction information between receptors and peptides from evolutionary information. In addition, our results indicate that AF2 is not just a modeling tool for monomeric structures, but also can be an extremely powerful tool for modeling more complex quaternary structures of proteins.

## Notes

### Competing Interest Statement

The authors have declared no competing interest.

### Summary of Updates

Cited more references from twitter.

